# Platelet disturbances correlate with endothelial cell activation in uncomplicated *Plasmodium vivax* malaria

**DOI:** 10.1101/715003

**Authors:** João Conrado Khouri Dos-Santos, João Luiz Silva-Filho, Carla C Judice, Ana Carolina Andrade Vitor Kayano, Júlio Aliberti, Ricardo Khouri, Diógenes S. de Lima, Helder Nakaya, Marcus Vinicius Guimarães Lacerda, Erich Vinicius De Paula, Stefanie Costa Pinto Lopes, Fabio Trindade Maranhão Costa

## Abstract

**Introduction:** Platelets drive endothelial cell activation in many diseases. However, if this occurs in *Plasmodium vivax* malaria is unclear. As platelets have been reported to be activated and to play a role in inflammatory response during malaria, we hypothesized that this would correlate with endothelial alterations during acute illness.

**Methods:** We performed platelet flow cytometry of PAC-1 and P-selectin. We measured Platelet markers (CXCL4, CD40L, P-selectin, Thrombopoietin, IL-11) and endothelial markers (ICAM-1, von Willebrand Factor and E-selectin) in plasma with a multiplex-based assay. The values of each mediator were used to generate heatmaps, K-means clustering and Principal Component analysis. In addition, we determined pair-wise Pearson’s correlation coefficients to generate correlation networks.

**Results:** Platelet counts were reduced, and mean platelet volume increased in malaria patients. The activation of circulating platelets in flow cytometry did not differ between patients and controls. CD40L levels (Median [IQ]: 517 [406-651] vs. 1029 [732- 1267] pg/mL, *P*=0.0001) were significantly higher in patients, while P-selectin (Median [IQ]: 17.0 [15.4-20.6] vs. 22.2 [17.6-25.7] ng/mL, *P*=0.0621) and CXCL4 showed a nonsignificant trend towards higher levels in patients. The network correlation approach demonstrated the correlation between markers of platelet and endothelial activation, and the heatmaps revealed a distinct pattern of activation in two subsets of *P. vivax* patients when compared to controls.

**Conclusion:** platelet activation occurs in uncomplicated vivax malaria and this correlates with higher endothelial cell activation, especially in a subset of patients.

**AUTHOR SUMMARY:** Endothelial cell activation is a key process in the pathogenesis of *Plasmodium vivax* malaria. Platelets are classically involved endothelial cell activation in several diseases, but their role in context of vivax malaria remains unclear. Thrombocytopenia is the most common hematological disturbance in *P. vivax*-infected patients, and platelets have been implicated in parasitemia control. In this study, we studied the activation of platelets in association with endothelial cell activation in vivax malaria. Platelets retrieved from infected peripheral blood were non-activated when analyzed by flow cytometry; however, they displayed higher mean volume and significantly reduced counts. We also found higher levels of circulating factors associated with platelet activation (especially CD40L), thrombopoiesis and endothelial cell activation in infected patients. Further, through pair-wise correlation and clustering analysis, we found a subgroup of patients showing significant associations between markers of platelet and endothelial activation in a pattern different from that of endemic controls. Collectively, our findings point to a peculiar role of platelets in endothelial cell activation in vivax malaria and indicate a heterogeneous host response in the setting of uncomplicated disease, a finding to be further explored in future studies.

## INTRODUCTION

Thrombocytopenia is the most common hematological alteration in malaria, although there is no definitive mechanistic explanation to its occurrence (1, 2). In a retrospective cohort, patients who died from malaria had lower platelet counts in comparison to those with less severe disease (3). Moreover, *P. vivax* malaria patients with thrombocytopenia showed higher levels of markers of endothelial cell (EC) activation compared to those with normal platelet counts (4) and some grade of platelet activation has been reported during the disease (5). Together, these reports point out to a significant role of platelets in the pathophysiology of malaria.

In cerebral malaria, the most severe presentation of the disease, platelets have been shown to accumulate in the brain microvessels of affected children (6). In mice, the adhesion of platelets to brain endothelial cells was crucial for the development of the syndrome (7). In addition, platelets lead to a deleterious inflammatory response in the disease through platelet-factor 4 (PF4), with PF4-KO mice surviving the infection (8). However, PF4 also plays a protective role in the disease, as it mediates *P. falciparum* killing by platelets *in vitro* (9) and *in vivo*, as shown in patients from Southeast Asia, and this was correlated with reduced parasitemias (10).

EC activation is present in all malaria species, occurring in both mild and severe cases (11). In terms of pathogenesis, EC activation is important for *P. falciparum*-infected erythrocytes adhesion to microvasculature, avoiding immunological clearance and leading to severe disease, while inducing more EC damage (12). *P. vivax*-infected erythrocytes also adhere to EC (13), but the magnitude of the phenomenon is smaller and if this has a role in endothelium pathology and disease severity is not clear (14).

While the role of platelets in *P. falciparum* malaria has been extensively studied, its role in the pathogenesis of EC dysfunction during *P. vivax* malaria remains to be investigated. In this study, we show that platelet counts were reduced in *P. vivax* malaria patients, while circulating markers of platelet activation were higher. Importantly, platelet activation markers correlated with those related to endothelial activation, indicating a role for platelets in EC pathology in this disease.

## METHODS

### Ethics Statement

All subjects enrolled in the study were adults, and samples were taken only after signing of informed consent. The study was approved by the local Research Ethics Committee at Fundação de Medicina Tropical Dr. Heitor Vieira Dourado (FMT-HVD, Manaus, Brazil), under #CAAE: 54234216.1.0000.0005. Sixty-five patients with *P. vivax* malaria, as diagnosed by light microscopy, seen at FMT-HVD and 37 healthy controls were enrolled. Exclusion criteria: under 18 years of age; pregnancy; in use of antimalarials; chronic disease; medication known to interfere with platelet count/function; smoking. After signing the informed consent, 20 mL of venous blood were drawn by venipuncture in a syringe with 15% acid citrate dextrose as anticoagulant to minimize in vitro platelet activation.

### Platelet isolation and poor platelet plasma preparation

Whole blood was centrifuged at 180 g for 18 minutes at room temperature, without brake for gradient formation, to obtain the platelet rich plasma (PRP). The PRP was centrifuged at 100 g for 10 minutes for removal of residual leukocytes, and subsequently centrifuged at 800g for 20 minutes to obtain the platelet pellet, prostaglandin E1 300 nM was used to minimize platelet aggregation. The supernatant of this centrifugation was centrifuged at 1000 g for 10 minutes to obtain platelet poor plasma (PPP).

### Platelet parameters

Within 15 minutes of sampling, complete blood counts were performed. Platelet activation was assessed in PRP using anti-CD61 antibody, and PAC-1 (FITC) and anti-P-selectin (PE) antibodies, by flow cytometry (FACSCanto, BD) and analysis with FlowJo software (Free Star). The same panel was used to assess whether the incubation of PPP (50% v/v for 10 min at 37 ºC) from malaria patients was capable to activate platelets from a healthy donor.

We also measured circulating factors associated with platelet activation and production in the patients’ plasma, using a multiplex-based cytokine assay (R&D Systems): CD40L, P-selectin, PF4 and thrombopoietin (TPO), IL-11, as well as circulating markers of EC activation (ICAM-1, E-selectin, von Willebrand Factor (vWF)). We selected 31 patients for the multiplex assay from a wide range of parasitemias (260 to 25,150 *Pv*-IE/μL), so that a wide spectrum of disease burden could be evaluated. This subgroup did not differ from the overall population of patients regarding severity of disease, sex proportion, platelet counts and mean parasitemia. We selected nine controls matched for age and sex. For the network analysis, we used some interpolated results from below standard range.

### Network and clustering analysis

The values of each circulating factor measured in the plasma samples, hematological parameters and parasitemia from endemic controls and *P. vivax* malaria patients were input in the R software (v 3.4.3) to generate heatmaps and to perform K-means clustering. After running the algorithms, individuals were clustered according to the levels of expression of the mediators in 3 groups, which were named Control, Vivax^low^ and Vivax^high^. In addition, the same software was used to determine pair-wise Pearson’s correlation coefficients to generate correlation networks and the p value to test for non-correlation was evaluated using p ≤ 0.05 as a cut-off. In order to analyze the structure of the networks, the graphics for the network analysis were customized in the Cytoscape software (v 3.5.1) using the prefuse force-directed layout, which in the equilibrium state for the system of forces, determined by the correlation strength, the edges tend to have uniform length, and nodes that are not connected by an edge tend to be drawn further apart.

### Statistical Analysis

Fisher’s exact test was used for categorical data. Student’s t-test was used to compare means between groups with normally distributed data, and data sets with non-normal distributions were compared using the Mann–Whitney test, with p<0.05 considered significant. Data are presented as means and SD unless otherwise stated. Analysis were performed, and the graphs generated in GraphPad Prism5 and R software.

## RESULTS

### Platelet parameters in malaria patients

Platelet counts were significantly reduced in patients (Mean ± SD: 91.5±41.3 ×10^9^/L vs. 244.1±52.1 ×10^9^/L, P<0.0001), yielding an 88,8% frequency of thrombocytopenia (Figure 1A). Mean Platelet Volume (MPV) was increased in patients (9.2±1.1 fL) compared to controls (8.7±0.6 fL, P<0.0126) (Figure 1B), and was inversely correlated with platelet counts in both patients and control groups (Patients r= −0.4959, Controls r= −0.6898, both P<0.0001) (Figure 1C). Platelet counts were not correlated with parasitemia.

**Figure 1.**
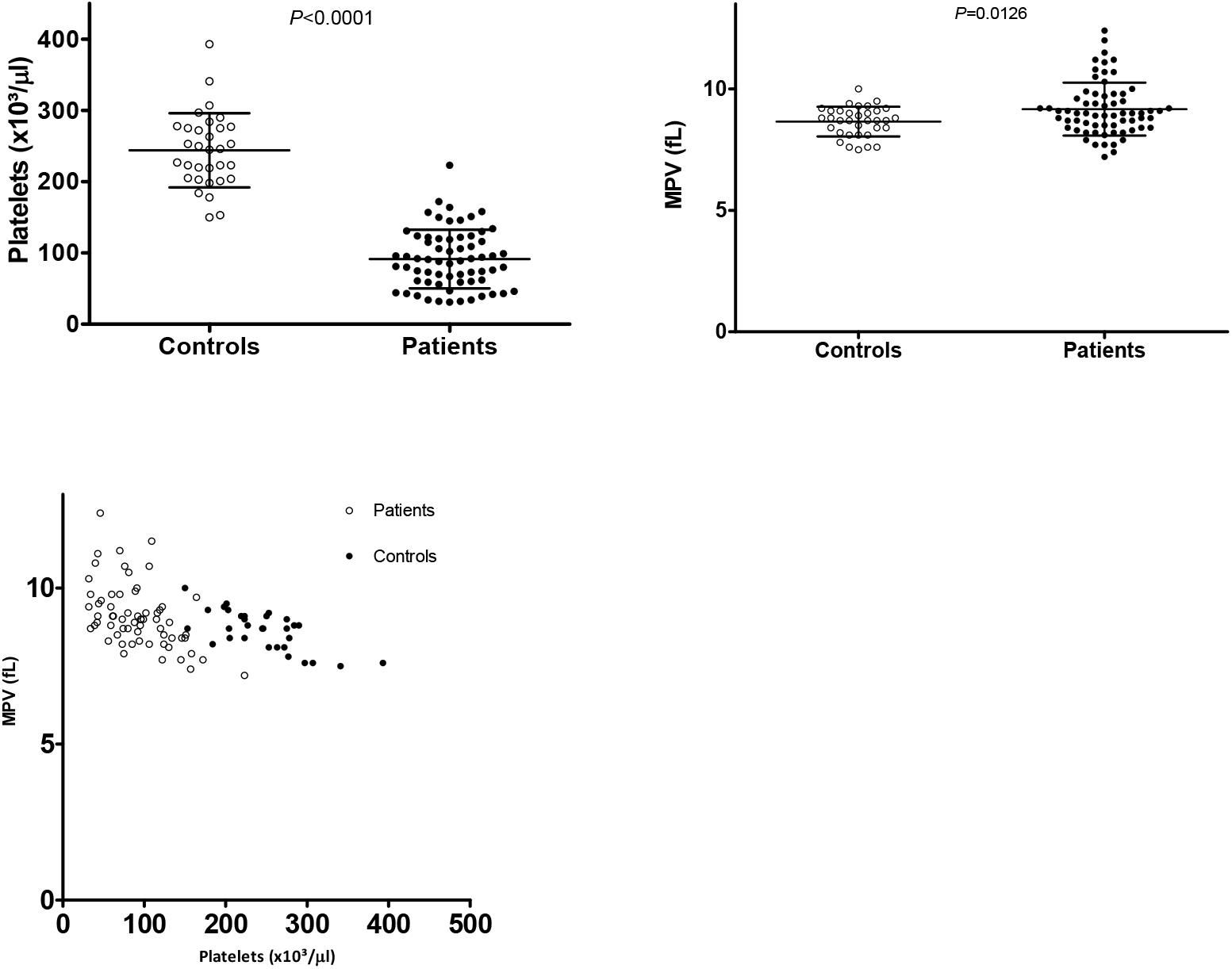
Platelet counts and volume. **A)** Platelet counts were significantly reduced in malaria patients. **B)** MPV was moderately increased in patients. **C)** Platelet count and MPV were inversely correlated in both patients and controls.

### Platelet Activation in vivax malaria

Platelet activation is a feature of some thrombocytopenic infections as well of diseases associated to endothelial cell dysfunction (15, 16). There was no significant difference in the percentage of expression of P-selectin and PAC-1 platelets between patients and controls in flow cytometry (Figure 2A-B).

**Figure 2.**
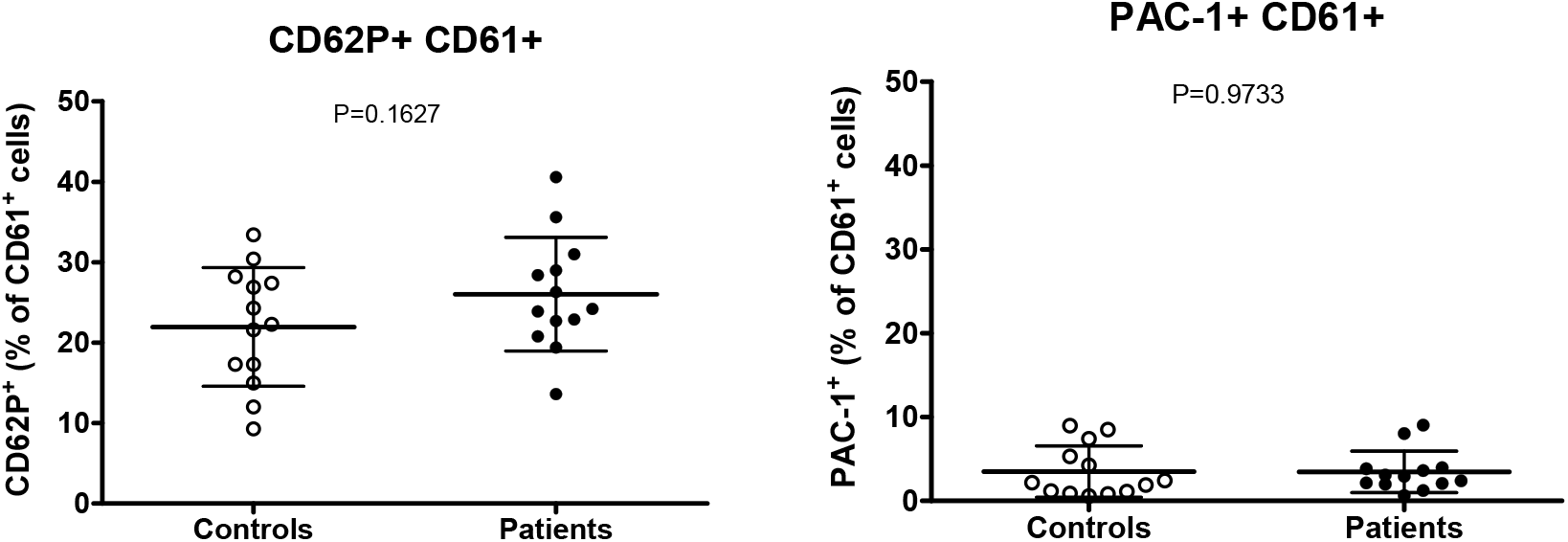
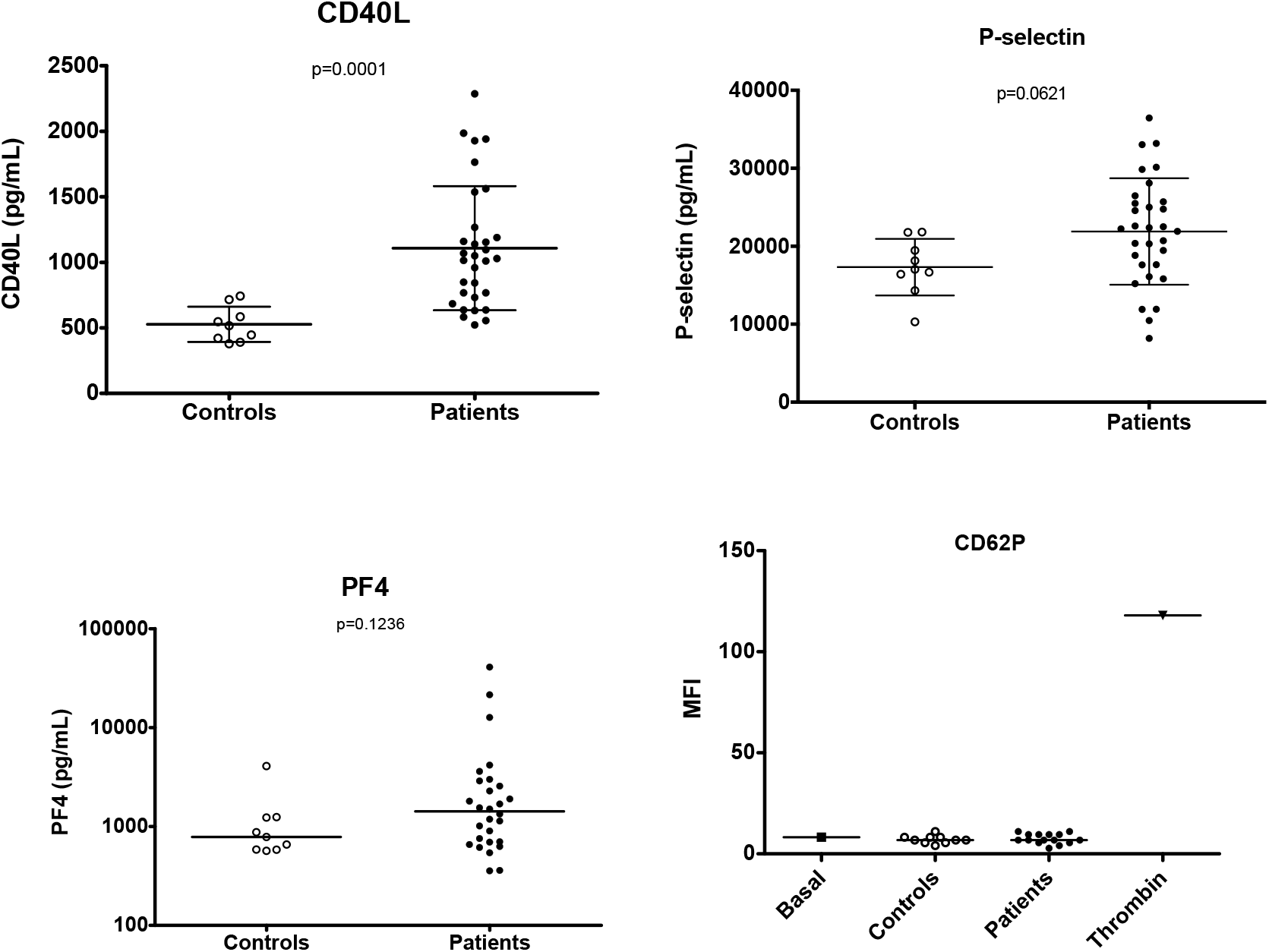
Platelet activation. Percentage of platelet activation by **A)** anti-P-selectin and **B)** PAC-1 staining (N=11). **C)** Levels of sCD40L. **D)** Levels of sPselectin. **E)** Levels of PF4 (Controls, n=9, Patients n=31). **F)** Plasma form malaria patients failed to induce activation of controls platelets. We used thrombin as a positive control

In contrast, patients had higher levels of CD40L in plasma (Median [IQ]: 517 [406-651] vs. 1029 [732-1267] pg/mL, *P*=0.0001), with a 2-fold increase in comparison to controls. P-selectin showed a trend towards elevated levels in patients (Median [IQ]: 17.0 [15.4-20.6] vs. 22.2 [17.6-25.7] ng/mL, *P*=0.0621), while PF4 levels (Median [IQ]: 784 [584-1239] vs. 1420 [683-2813] ng/mL, *P*=0.1236) were not different between the groups (Figure 2C-E, Table 1). Patients’ PPP failed to activate platelets in comparison to PPP from controls (Figure 2F).

**Table 1.** Plasma levels of markers of platelet activation and production.

| Parameter | Controls (n=9) | Patient (n=31) | P |
| --- | --- | --- | --- |
| Median [IQ 25-75] |  |  |  |
| **CD40L (pg/mL)** | 517 [406-651] | 1029 [732-1267] | 0.0001 |
| **P-Selectin (ng/mL)** | 17.0 [15.4-20.6] | 22.2 [17.6-25.7] | 0.0621 |
| **PF4 (pg/mL)** | 784 [584-1239] | 1420 [683-2813] | 0.1236 |
| **TPO (pg/mL)** | 2046 [1652-2194] | 2996 [2554-3402] | <0.0001 |
| **IL-11 (pg/mL)** | 3547 [2904-4298] | 5748 [4687-6448] | <0.0001 |

### Thrombopoiesis

Acute reductions in platelet numbers and inflammatory states disturb thrombopoiesis (17, 18). Therefore, we measured the circulating levels of the cytokines thrombopoietin (TPO) and IL-11, important players in the production of platelets in health and disease. An 50% increase in these two markers was observed in malaria patients (Table 1), and they were significantly correlated (Pearson r = 0.8476, 95%CI: 0.7049-0.9243, P<0.0001). There was no correlation between TPO and platelet counts (Figure 3).

**Figure 3.**
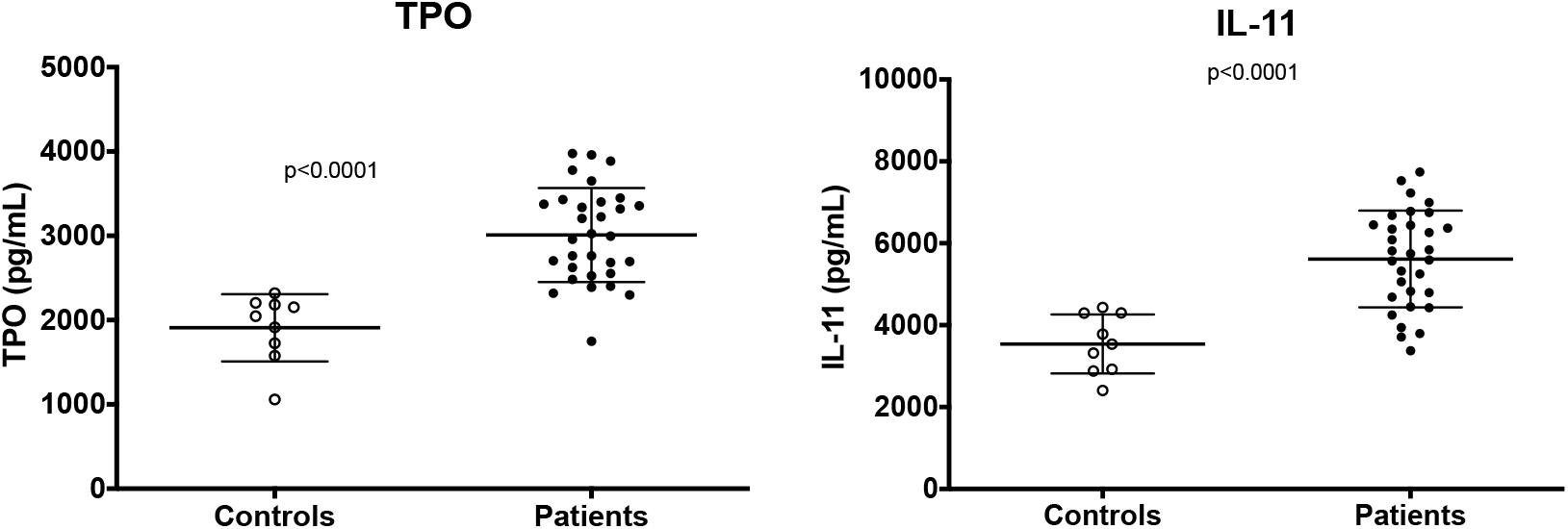
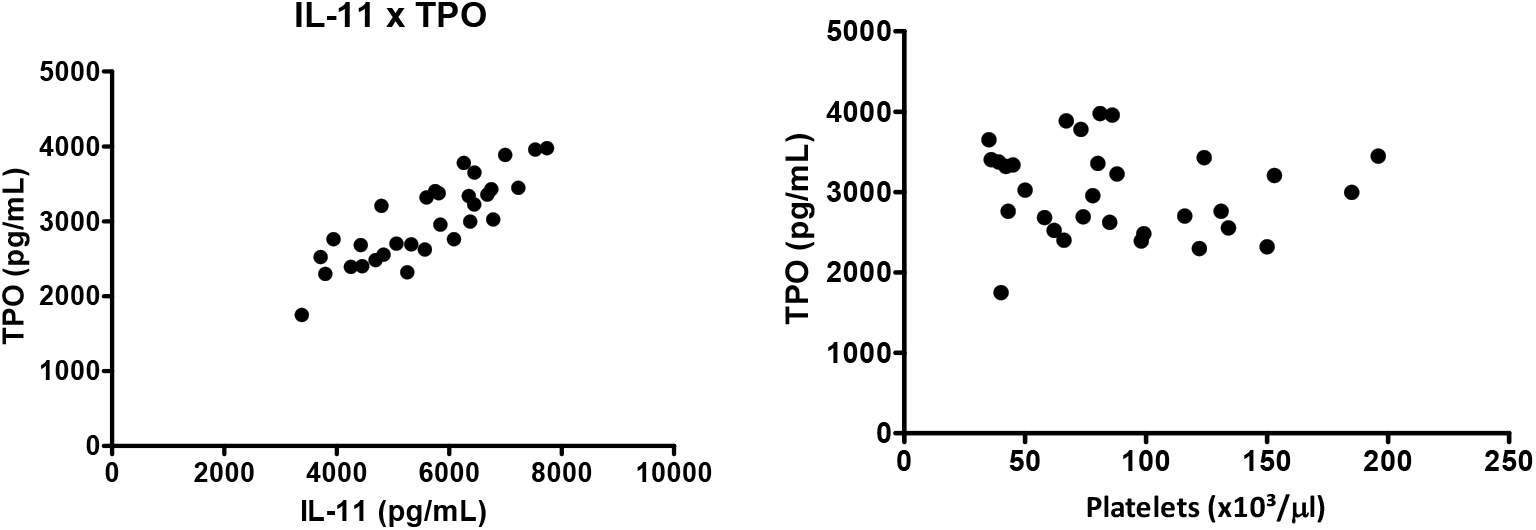
Levels of thrombopoietic cytokines in*P. vivax* malaria. **A)** Elevated levels of TPO in patients compared to controls. **B)** Elevated IL-11 in patients compared to controls. **C)** Significant association between TPO and IL-11 (r=0.8476, 95%CI: 0.7049-0.9243, P<0.0001), indicating a concerted stimulus to platelet production. **D)** Lack of association between TPO and platelet counts, indicating that the “sponge model” does not solely explain regulatory mechanisms of TPO production in this disease.

### Correlations and networks

Markers of platelet activation, thrombopoiesis and EC activation were significantly higher in *P. vivax* malaria patients in relation to endemic controls. Importantly, all the associations between platelet and EC markers were positive, a change of pattern in relation to controls, in which both negative and positive associations occurred (Figure 4A-B). Parasitemia was significantly correlated with the markers of thrombopoiesis TPO and IL-11 and with ICAM-1 (Figure 4B).

**Figure 4.**
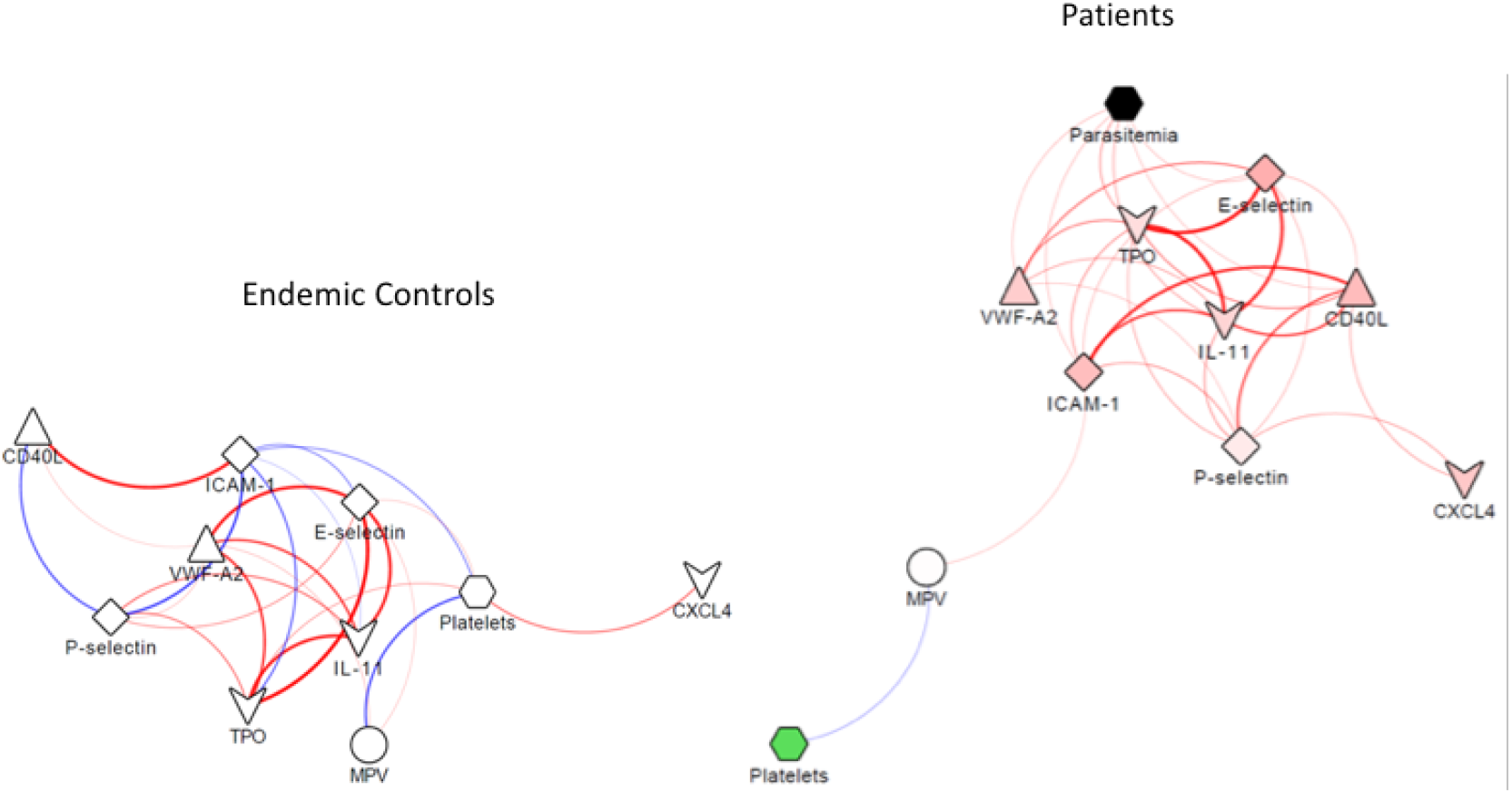
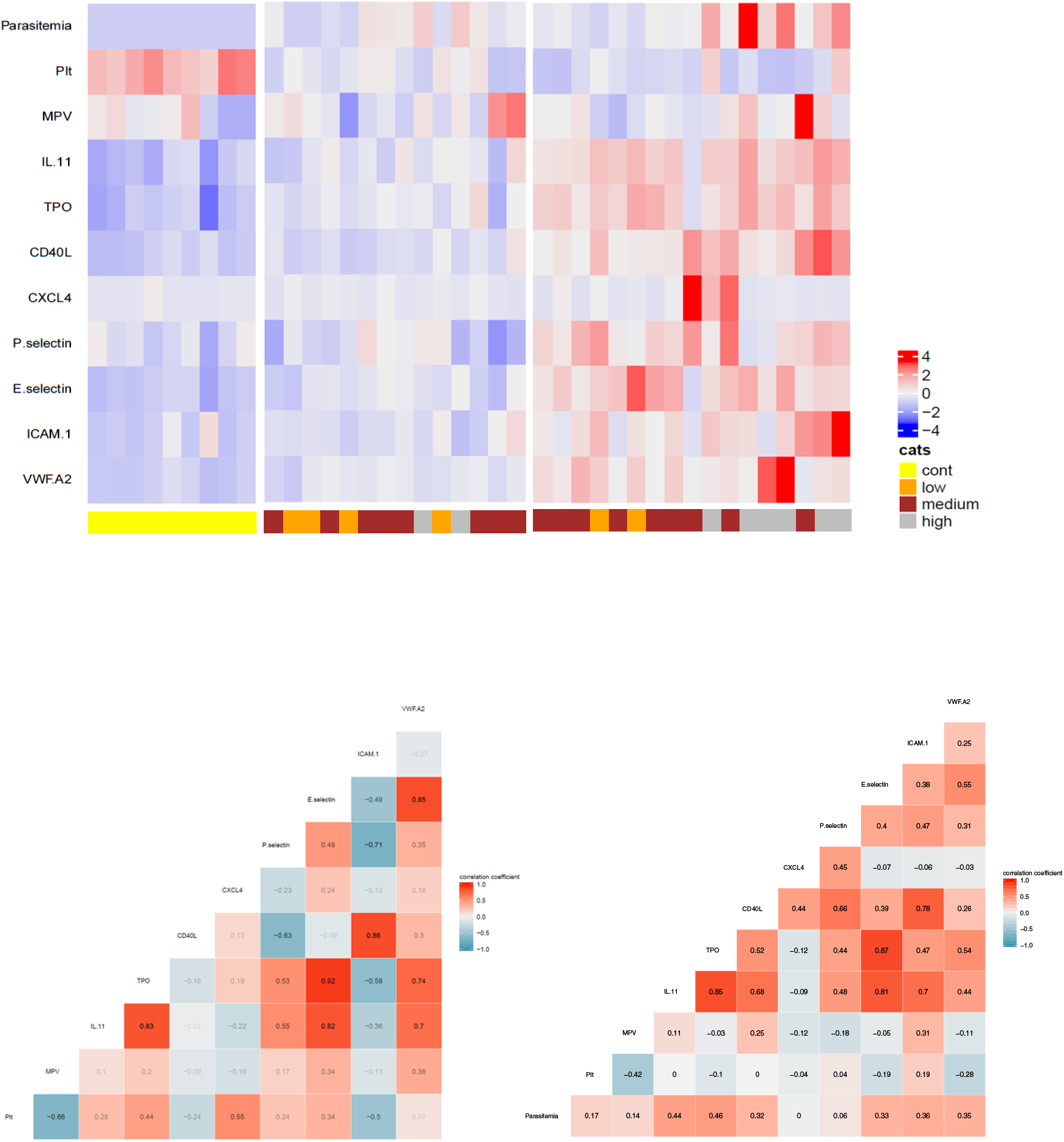
Correlation between markers of platelet and endothelial cell activation. Networks of the correlations between platelet parameters and markers of endothelial cell activation in controls **(A)** and patients **(B)**, using a prefuse force-directed layout done in Cytoscape software (v 3.5.1). **C)** Heatmap obtained from the normalized expression of parasitemia, platelet parameters and markers of endothelial cell activation, showing three distinct clusters of controls and two groups of patients. Yellow: Controls; Orange: low parasitemia; Brown: moderate parasitemia; Gray: high parasitemia. Value for each pair of correlation between the parameters measured in the plasma from controls **(D)** and patients **(E)**.

Additionally, the heatmap generated from the expression of the analytes revealed a defined separation of controls and two different groups of *P. vivax* malaria patients (Figure 4C). In comparison to endemic control individuals one group of *P. vivax* malaria patients had a lower overall variation in the response. The second group of patients clearly showed a more potent response to the infection, with a higher variation in the expression of platelet activation and EC activation markers. Figure 4D-E displays the value of each pair of correlations.

## DISCUSSION

In the current study, we aimed to assess platelet activation and its relationship with endothelial cell activation in the context of *P. vivax* malaria. Thrombocytopenia was the most frequent hematological alteration in our cohort and, as previously reported for malaria (1), was correlated to an increase in MPV. Several disease states in which platelets are consumed mainly in the periphery present with the same correlation. The slightly increased MPV in malaria patients could be important during the disease as larger platelets are thought to be more active in comparison to smaller ones, and have been implicated in diseases were endothelial activation and dysfunction play a central role (16). However, in this study, MPV showed only a non-significant correlation with ICAM-1 levels.

Adding to the investigation of platelet disturbances during the disease, we also measured the levels of TPO and IL-11, two cytokines central to platelet production. Previous reports have demonstrated a high platelet turnover during malaria (19) and increased in megakaryocyte numbers in bone marrow (20). The observed rise in TPO and IL-11 levels in our cohort indicates that increased platelet production occurs during vivax malaria in response to the reduction in circulating platelet counts. Nonetheless, TPO levels in *P. vivax* patients presented no correlation with platelet counts, as opposed to what would be expected based on the classical “sponge” model – whereby TPO levels are regulated simply by platelet number – of the regulation of thrombopoiesis. These results indicate that during *P. vivax* malaria, thrombopoiesis might be regulated by additional mechanisms known to stimulate TPO production, such as IL-6 or activation of the Ashwell-Morell receptor (21). Interestingly, TPO and IL-11 were positively associated with the markers of platelet and EC activation, highlighting the correlation of thrombopoiesis alterations and inflammatory states (16). Whether these alterations in platelet production in an acute disease as malaria have major implications for platelet function – as has been shown for chronic inflammatory diseases (15) – is a relevant question for future studies.

Whether a systemic activation of circulating platelets occur in malaria is still unclear. While a study has reported altered platelet responses after exposure to *P. falciparum* infected erythrocytes (P-IE) *in vitro* (22), direct assessment of platelet activation through flow cytometry has rendered negative results (10, 23). However, evidence of some grade of platelet activation *in vivo*, through measurement of circulating factors, have been shown by other groups (5, 24). Therefore, on the one hand, our flow cytometry results, both in platelets from patients and in platelet stimulation with patients’ plasma, are in line with has been previously reported in the literature. On the other hand, our results of a trend towards elevated (albeit not significantly) P-selectin and, especially, elevated CD40L, argue for some grade of systemic platelet activation, leading to platelet degranulation. Interestingly, platelets have been shown to release bioactive sCD40L during vaso-oclusive in sickle cell disease, a finding that implicates platelet-derived sCD40L in vascular events (25). Moreover, incubation of activated platelets with cultured endothelium leads to its activation through CD40L-CD40 interaction (26). However, as CD40L is also a molecule involved in T-cell activation, we cannot exclude that the latter is a relevant source of the sCD40L found in the patients.

Endothelial cell activation is a major component of malaria pathogenesis (11), a phenomenon with an extensive participation of platelets in different disease settings (16). As platelets are classically associated with endothelial pathology, we searched for patterns of association between platelet factors and markers of endothelial cell activation. Interestingly, our networks revealed the association of soluble CD40L and P-selectin with ICAM-1 and E-selectin, indicating the interplay between these two cell populations in the disease, with a possible role for platelets in the pathogenesis of endothelial cell activation in vivax malaria. CD40L has been described as a central molecule in the pathogenesis of atherosclerosis and is elevated during vaso-oclusive crisis in sickle cell disease (25), features that argue for a potential causative role of this molecule in EC activation in malaria.

Notably, the heatmaps further confirmed a distinct pattern of platelet and endothelial cell activation in a subset of patients (Figure 2), which were more markedly distinct from controls. This finding reinforces the hypothesis that platelet activation and release of granule content plays a role in the endothelium alterations in *P. vivax* malaria (5).

A limited small number of patients were included in the multiplex-based assay, limiting the generalizability of the findings and the statistical analyses. Moreover, while the role for platelets in endothelial activation in malaria is indicated by our results and has biological foundation, the mechanisms behind this association were not elucidated. Finally, although all patients included in the study presented with uncomplicated malaria, we did not follow patients throughout their illness to assess potential clinical implications of our findings.

In this study, we found evidence of platelet activation during vivax malaria. Importantly, we show that soluble CD40L is elevated during the infection and the positive associations with markers of EC activation indicate that this molecule is a player in the pathogenesis of this disease. Therefore, we believe that future studies to identify the mechanisms of how platelets induce EC pathology in relevant models of vivax malaria are warranted.

## Acknowledgements

We would like to thank the support of the malaria diagnosis team at Fundação de Medicina Tropical. Cytometric analysis was done at the Flow Cytometry Platform at Instituto Leônidas & Maria Deane, Fiocruz Amazônia. The multiplex was performed at the Laboratório Central de Tecnologias de Alto Desempenho (LaCTAD, University of Campinas).

## Meetings

Some of the results from this paper have been previously presented at the Brazilian Congress of the Hematology and Hemotherapy (2018) and the International Conference of Plasmodium vivax Research (2019) in the form of posters.

